# A special satellite-like dsRNA of a novel hypovirus from *Pestalotiopsis fici* broadens the definition of satellite

**DOI:** 10.1101/2022.09.22.508974

**Authors:** Zhenhao Han, Jiwen Liu, Linghong Kong, Yunqiang He, Hongqu Wu, Wenxing Xu

**Author notes:** These authors contributed equally to this work (Zhenhao Han, Jiwen Liu). Corresponding author Wenxing Xu, Professor, Plant Pathology, Hongqu Wu, PhD, Forest Pathology.

## Abstract

Satellites associated with plant or animal viruses have been largely detected and characterized, while those from mycoviruses together with their roles remain far less determined. Three dsRNA segments (dsRNA 1 to 3 termed according to their decreasing sizes) were identified in a strain of phytopathogenic fungus *Pestalotiopsis fici* AH1-1 isolated from a tea leaf. The complete sequences of dsRNAs 1 to 3, with the sizes of 10316, 5511, and 631 bp, were determined by random cloning together with a RACE protocol. Sequence analyses support that dsRNA1 is a genome of a novel hypovirus belonging to a newly proposed genus “Alphahypovirus” of the family *Hypoviridae*, tentatively named Pestalotiopsis fici hypovirus 1 (PfHV1); dsRNA2 is a defective RNA (D-RNA) generating from dsRNA1 with septal deletions; and dsRNA3 is the satellite component of PfHV1 since it could be co-precipitated with other dsRNA components in the same sucrose fraction by ultra-centrifuge, suggesting that it is encapsulated together with PfHV1 genomic dsRNAs. Moreover, dsRNA3 shares an identical stretch (170 bp) with dsRNAs 1 and 2 at their 5′ termini and the remaining is heterogenous, which is distinct from a typical satellite that generally has very little or no sequence similarity with helper viruses. More importantly, dsRNA3 lacks a substantial open reading frame (ORF) and a poly (A) tail, which is unlike the known satellite dsRNAs of hypoviruses, as well as unlike those in association with *Totiviridae* and *Partitiviridae* since the latters are encapsidated in coat proteins. As up-regulated expression of dsRNA3, dsRNA1 was significantly down-regulated, suggesting that dsRNA3 negatively regulates the expression of dsRNA1, whereas dsRNAs 1 to 3 have no obvious impact on the biological traits of the host fungus including morphologies and virulence. This study indicates that PfHV1 dsRNA3 is a special type of satellite-like nucleic acid that has substantial sequence homology with the host viral genome without encapsidation in a coat protein, which broadens the definition of satellite.

**IMPORTANCE:** Satellites in association with plant or animal viruses have been largely detected and characterized, while those from mycoviruses together with their roles remain far less determined. Here, a special satellite-like dsRNA (SatL-dsRNA) together with its helper virus, a novel hypovirus from *Pestalotiopsis fici*, was identified and characterized. This SatL-dsRNA lacks a substantial open reading frame and a poly (A) tail, which is unlike the known satellite dsRNAs of hypoviruses. It is also unlike those in association with *Totiviridae* and *Partitiviridae* since the latters are encapsidated in coat proteins. As up-regulated expression of the SatL-dsRNA, the helper virus genome was significantly down-regulated, suggesting that it negatively regulates the genomic expression of the helper virus. This special SatL-dsRNA has substantial sequence homology with the host viral genome and is not encapsidated in the coat protein of its helper virus, which represents a novel class of satellite-like nucleic acids, and it broadens the definition of satellite.

## INTRODUCTION

*Hypoviridae* is a family of capsidless viruses with positive-sense (+), single-stranded (ss) RNA genomes of 9.1–12.7 kb that possess either a single large open reading frame (ORF) or another small ORF (1). Only one genus, *Hypovirus*, is included in this family, which contains Cryphonectria hypovirus 1-4 (CHV1-4) and is associated with biologic control of the filamentous fungus that causes chestnut blight (2). With the increase in the number of identified hypoviruses, three proposed genera have been proposed, namely “Alphahypovirus”, “Betahypovirus” and “Gammahypovirus” (3, 4). Hypoviruses have been heavily detected in ascomycetous and basidiomycetous filamentous fungi, and some of these were able to alter fungal host phenotypes and attenuate the host virulence, e.g., Alternaria alternata hypovirus 1 (AaHV1)(5), Botrytis cinerea hypovirus 1 (BcHV1)(6), while others were not, e.g., Fusarium graminearum hypovirus 1 (FgHV1)(7). Defective RNAs (D-RNAs) were generally detected in association with hypoviruses, e.g., CHV1-EP713 (8, 9), Sclerotinia sclerotiorum hypovirus 1 (SsHV1)(10), Fusarium graminearum Hypovirus 2 (FgHV2)(11), Botrytis cinerea hypovirus 1 (BcHV1)(6), AaHV1 (5), which were a type of generations of internally deleted genomic RNAs.

Satellites are subviral agents which lack genes that could encode functions needed for replication, and depend on the co-infection of a host cell with a helper virus, which consists of satellite viruses and satellite nucleic acids, discriminating by the presence of a structural protein (12). Satellite nucleic acids include ssDNAs and ssRNAs, associated with plant or animal viruses, and double stranded RNAs (dsRNAs), are mainly associated with mycoviruses, namely the families *Totiviridae* and *Partitiviridae*. Satellites do not constitute a homogeneous taxonomic group, and have no taxonomic correlation with their helper viruses, and have arisen independently a number of times during virus evolution. Since satellites were mostly characterized as ssRNA satellites that use ssRNA plant viruses as helpers, and satellite dsRNAs remain very limited characterized and it is very likely that other satellites, some with novel combinations of characters, remain to be discovered.

*Pestalotiopsis* spp. belongs to the family Amphisphaeriaceae that are distributed widely throughout tropical and temperate regions. They are common phytopathogens reducing production and causing enormous losses in either horticultural plants or forest trees (13). For example, in tea plants, *P. theae* and *P. longiseta* are responsible for tea gray blight disease, which leads to large brown spots with the formation of apparent concentric rings in the late period of disease on mature and old leaves of tea plants, and even leads to death of tender branches of infected tea seedlings; *P. theae, P. camelliae, and P. clavispora* were also identified in association with small brown-black spots on tea tender leaves (14, 15). However, endophytic *Pestalotiopsis* species can confer fitness to the host plants, and are also thought to be a rich source for bioprospecting compared to other fungal genera since they produce novel compounds with medicinal, agricultural and industrial applications (15). Therefore, *Pestalotiopsis* fungi live in host plants with a complex lifestyles, being parasitic, commensal, or mutualistic. Mycoviruses might have participated in the transition of these lifestyles, since a chrysovirus, the first mycovirus isolated from *Pestalotiopsis* fungi, can convert its host fungus from a phytopathogenetic one to a nonpathogenetic endophyte on tea leaves, conferring high resistance to the host plants against the virulent *P. theae* strains (13). It is fascinating to characterize more mycoviruses to understand the virus taxonomy, evolution, molecular and biological traits related to this important fungal species. Specifically, characterization of a hypovirus in this fungal species remains a great interesting since hypoviruses represent the classic and successful examples for biological control of plant diseases (16).

In this study, a novel hypovirus from *Pestalotiopsis fici* together with a novel special satellite dsRNA, were identified and characterized, which represents the first hypovirus and the second mycovirus isolated from *Pestalotiopsis* spp., as well as a novel class of satellite nucleic acids, illustrating some novel molecular and biological traits of a hypovirus.

## MATERIALS AND METHODS

### Fungal strains and cultures

*P. fici* strain AH1-1 was isolated from a tea leaf showing the typical symptoms of tea grey blight disease collected in Liuan city, Anhui province, China, and was identified based on morphologies and multi-loci sequences as previously described (17), and strain AH1-1-14 is a subisolate generated from a single conidium of strain AH1-1. Strain AH1-1V^-^ was a cured subisolate of AH1-1 by elimination of dsRNA components, and strain AH1-1^hyg^V^-^ was generated from AH1-1V^-^ labeled with Hygromycin (Hyg). *P. theae* TP-2-2W, JWX-3-1, and CJB-4-1 were collected from tea leaves in Yichang city, Hubei province, China, and were identified on their morphologies and ITS sequence. All strains were grown at 25°C in the dark for 3-5 days on solid potato dextrose agar (PDA; 20% diced potatoes, 2% glucose, and 1.5% agar) unless otherwise stated.

### dsRNA extraction, purification, and enzymatic treatments

For dsRNA extraction, fungal mycelial plugs were inoculated onto sterilized cellophane disks on PDA plates at 25ºC in the dark for 4 to 5 days, and the mycelia were collected and subjected to dsRNA extraction as previously described (18). The resulting nucleic acids were treated with 2 U DNase I (New England Biolabs), 10 U S1 nuclease (Thermo Scientific) at 37°C for 1 h, or those without treatments were fractionated by electrophoresis on 1.2% agarose gels with Tris-acetate-EDTA (TAE) buffer and visualized by staining with ethidium bromide. Each of the three dsRNAs was excised, purified using a gel extraction kit (Qiagen, USA), dissolved in DEPC-treated water and stored at −80 °C until use.

The dsRNA preparation, BdRV1 dsRNAs (extracted from mycelia infected by BdRV1) (19), *in vitro* transcripts of cDNAs of peach latent mosaic viroid (PLMVd) (14) with its linearized pGEM-T plasmids harboring the cDNAs, and DNA controls generated from PCR products were treated with S1 nuclease or/and DNase I as indicated above, and with 200 ng/mL RNase A (Thermo Scientific) in 2×SSC (300 mM NaCl, 30 mM sodium citrate, pH 7.0) or 0.1×SSC as previously described (18).

### Cloning, sequencing, and sequence analysis

The sequences of the dsRNAs were determined by cloning and sequencing amplicons generated by reverse transcription and polymerase chain reaction (RT-PCR) using the random primers 05RACE-3RT and 05RACE-3 (Table S1) as previously described (17). The 5′- and 3′-terminal sequences of the dsRNAs were obtained by cloning and sequencing the RT-PCR amplicons generated using a standard RNA ligase mediated rapid amplification of cDNA ends (RLM-RACE) protocol, including the usage of PC3-T7loop and PC2 (Table S1). The oligonucleotide primers used for RLM-RACE were designed based on sequence information obtained from the randomly primed amplicons (20). At least three independent clones of each amplicon were sequenced in both directions, by Sangon Biotech Co., Ltd, Shanghai, China. Sequence similarity searches were performed using the BLASTn program for nucleic acids or BLASTp for putative proteins against the National Center for Biotechnology Information (NCBI) databases. Multiple alignments of nucleic and amino acid (aa) sequences were conducted using MEGA7 and MAFFT online (<u>MAFFT alignment and NJ / UPGMA phylogeny (cbrc.jp)</u>). The phylogenetic tree for RdRp sequences was constructed using MEGA 11 with the Maximum Likelihood method (21). Open reading frames (ORFs) were deduced using an ORF finder (https://www.ncbi.nlm.nih.gov/orffinder/). Conserved domains were predicted using Phyre2 (https://www.sbg.bio.ic.ac.uk/phyre2).

### RT-PCR detection, RNA blot analysis, and co-precipitate analysis

RT-PCR amplification was performed using a specific primer pair derived from the dsRNA 1 sequence (PfHV1-1F: 5′-TTCGATTTCAACGCCAGGTC-3′; PfHV1-1R: 5′-GCCGGGTCTATCGTCTTTTC-3′) (Table S1), generating a 307-bp fragment with an annealing temperature of 56°C in a PCR Thermal Cycler (Model PTC-100, MJ Research, USA).

Digoxigenin (DIG)-labeled riboprobes 1 (binding to positions from 1588 to 1934 nt of dsRNA1), 2 and 3 (116-355 and 177-485 nt of dsRNA3, respectively) were synthesized *in vitro* transcription based on the corresponding cDNAs inserted into the pGEM-T easy vector with T7 RNA polymerase (Takara, Beijing, China) as previously described (22). Nucleic acids were spotted (for dot blotting) or electro-transferred (for Northern blot) to positively-charged nylon membranes (Roche Diagnostics), and hybridized with DIG-labeled riboprobes as previously described (22).

Co-precipitate analysis of dsRNAs was conducted by sucrose gradient centrifugation according to the methods used for the purification of viral particles previously described (13). The resulted aliquots of each fraction (100 μL) were subjected to dsRNA extraction to monitor for the presence of viral dsRNAs.

### Engineering of RNA3-over-expressed strain and qRT-PCR quantitative analysis

To generate RNA3-over-expressed strains, AH1-1^SatL-OE^, cDNA of RNA3 was amplified and an overexpression vector were constructed with the products. The vector was then transfected to the strain AH1-1V^-^ via PEG4000 mediated protoplasts transfection as previously described (13).

The total RNAs from mycelia were extracted by Trizol agent and subjected to digestion by 5×gDNA digesture (Foregene, Chengdu, China) to remove the genomic DNAs, followed by cDNAs synthesized using 2× reverse transcriptase (Foregene, Chengdu, China) and amplified using 2×TSINGKE master qPCR mix (Tsingke, Beijing, China) using specific primers designed for quantitative analysis in CFX96 Real-Time PCR Detection System (Bio-Rad, USA). Amplification was conducted for cycles of 95℃ for 10 s, 56℃ for 10 s and 72℃ for 15 s after initial heating at 95℃ for 1 min. A partial fragment of the *Actin* gene was used to normalize the RNA samples for each qRT-PCR, and each treatment was conducted with three technical replicates.

### Contact cultures of *P. fici* isolates

Horizontal transmission of the virus was conducted as previously described (18). Briefly, both donor and recipient strains were cultured together on the same Petri dishes at 25°C for 7 days, and allowed to physically contact each other. Following contact, mycelial agar plugs were excised from the contact area of two strains and were sub-cultured onto fresh PDA plates. Seven independent donor-recipient pairs were assessed, and two mycelial agar plugs were selected from each pair for further analysis, resulting in a total of 14 isolates.

### Growth rate, morphology, and virulence assays

Fungal growth rates and morphologies were assessed as previously described (13). Three biological replicates for each strain were monitored and the results subjected to statistical analysis as described below. Fungal virulence was determined following inoculation of detached tea leaves (*C. sinensis* vars. Echa no.1 or Fuyun no.6) as previously described (13). At 4 dpi, lesions that developed on the inoculated leaves were measured. Eight biological replicates for each strain were monitored and the results subjected to statistical analysis as described below.

### Statistical analyses

For the growth rate and virulence assays, the mean values for the biological replicates are presented as column charts with error bars representing standard error of mean (SEM). The graphs were produced in Excel (Microsoft) and GraphPad Prism 7 (GraphPad software). Identification of potential outliers, independent-samples t test and one-way analysis of variance (ANOVA) were performed using IBM SPSS Statistics. In summary, the normality tests indicated that all data sets were well-modeled by a normal distribution, therefore independent-samples t test was performed to assess only two groups of data, and one-way ANOVA was performed to assess more than two groups of data. *P*-values < 0.05 were considered to indicate statistical significance, while *P*-values < 0.01, extremely significant.

## RESULTS

### A complex of dsRNA segments were extracted from *P. fici* strain AH1-1

Nucleic acid preparations enriched in dsRNA were obtained from *P. fici* strains AH1-1 and TP-2-2W isolated from tea leaves collected in China, and were subjected to digestion with DNase I and S1 nuclease and then agarose gel electrophoresis. The results showed that three dsRNAs (nominated 1–3 according to their decreasing sizes) were detected in preparations of strain AH1-1 but not in a control strain TP-2-2W (Fig. 1B). Of these components, dsRNA1 migrated together with the fungal genomic DNAs as electrophoresis on an agarose gel, and became apparent after treatment by DNase I (Fig. 1B). The sequences of the full-length cDNAs of dsRNAs 1–3 were determined by assembling partial-length cDNAs that were amplified from the purified dsRNAs using RT-PCR and RLM-RACE protocols. The corresponding sequences were deposited in GenBank with accession numbers OP441373-OP441375.

**Figure 1.**
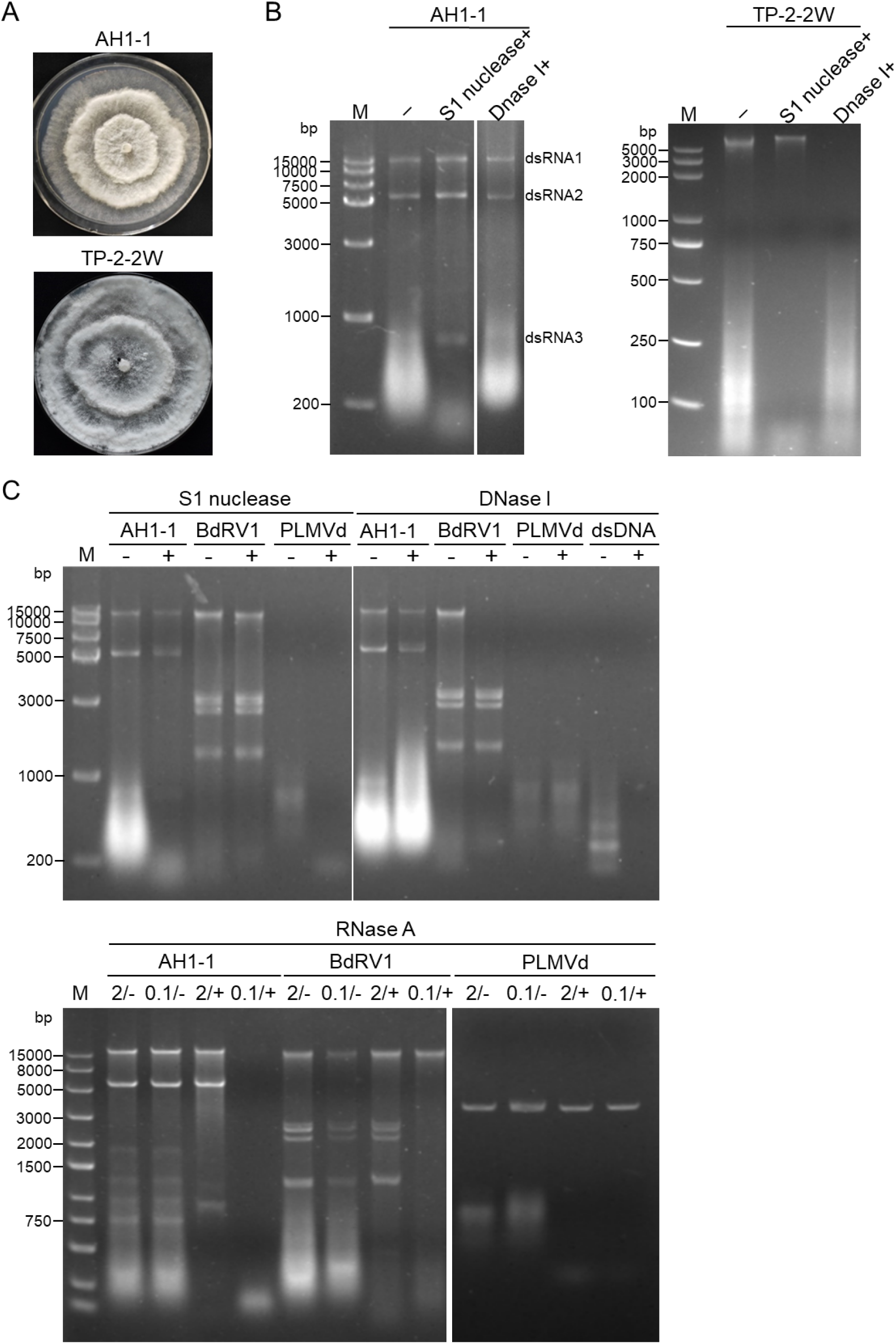
Fungal morphologies, and nucleic acid extraction and enzymatic treatments. (A) Colonies of *Pestalotiopsis* strain AH1-1 and TP-2-2W grown on PDA medium for 6 days. (B) Electrophoresis analysis of nucleic acids extracted form strain AH1-1 (left panel) and TP-2-2W (right) without treatment (lane 2) and treated with S1 nuclease (3) and DNase I (4), respectively, on 1% agarose gel. (C) Determined the dsRNA nature by enzymatic treatment with S1 nuclease, DNase I and RNase A. Botryosphaeria dothidea RNA virus 1 (BdRV 1), peach latent mosaic viroid (PLMVd), and PCR products were involved as dsRNA, ssRNA and DNA controls, respectively. M, DNA size marker.

The dsRNA nature of the three observed bands was further assessed by treatments with DNase III, S1 nuclease or RNase A (in 2× and 0.1×SSC), together with an ssRNA control (*in vitro* dimeric transcripts of peach latent mosaic viroid (PLMVd) and dsRNAs from a dsRNA mycovirus (Botryosphaeria dothidea RNA virus 1, BdRV 1). The RNAs extracted from strain AH1-1 together with BdRV 1 dsRNAs were digested by RNase A in 0.1×SSC, but they resisted digestion by DNase I, S1 nuclease, and RNase A in 2×SSC. In sharp contrast, PLMVd transcripts were completely degraded by S1 nuclease and by RNase A under both ionic conditions, but resisted digestion by DNase I; the genomic DNAs (extracted together with BdRV 1) and the cDNA plasmid of PLMVd (used for *in vitro* transcription) were completely degraded by DNase I while resistant to RNase A and S1 nuclease (Fig. 1C). These data strongly support that the nucleic acids extracted from strain AH1-1 were indeed dsRNAs instead of DNAs or ssRNAs.

### dsRNA 1 composes the genome of a novel hypovirus

The complete nucleotide sequence of dsRNA1 is 10316 base pair (bp) in length, consisting of a single putative open reading frame (ORF) beginning at AUG (nt positions 513-515) and terminating at UAG (9600-9602), coding for a polyprotein of 3029 aa with an approximate molecular mass of 345.9 kDa.

The deduced polyprotein aa sequence contains conserved domains of RNA dependent RNA polymerase (RdRp), viral RNA Helicase (Hel), and a most likely papain-like protease (Pro) (Fig. 2B). Within the RdRp domain, nine conserved motifs (Ia-VIII), characteristic for a RNA virus (23), were detected (Fig. 3A), and seven conserved motifs (I-VI) relatively conserved in (+) ssRNA were detected within the Hel domain (Fig. 3C); while in the possible Pro domain, only cysteine and glycine were detected whereas the core residue histidine was absent (Fig. 3B). Additionally, seven transmembrane domains (TMDs) were predicted among the entire polyprotein aa sequence (Fig. 2C). The 5′-untranslated region (UTR) of dsRNA1 is 512 nts in size, contains six AUG codons (Fig. S1C), and the 3′-UTR is 714 nts including a 6 nt-sized adenine tail (poly A) (Fig. 2B). Each UTR region forms a compact and complex stem-loop structure as predicted in RNAfold (Fig. S1A, B).

**Figure 2.**
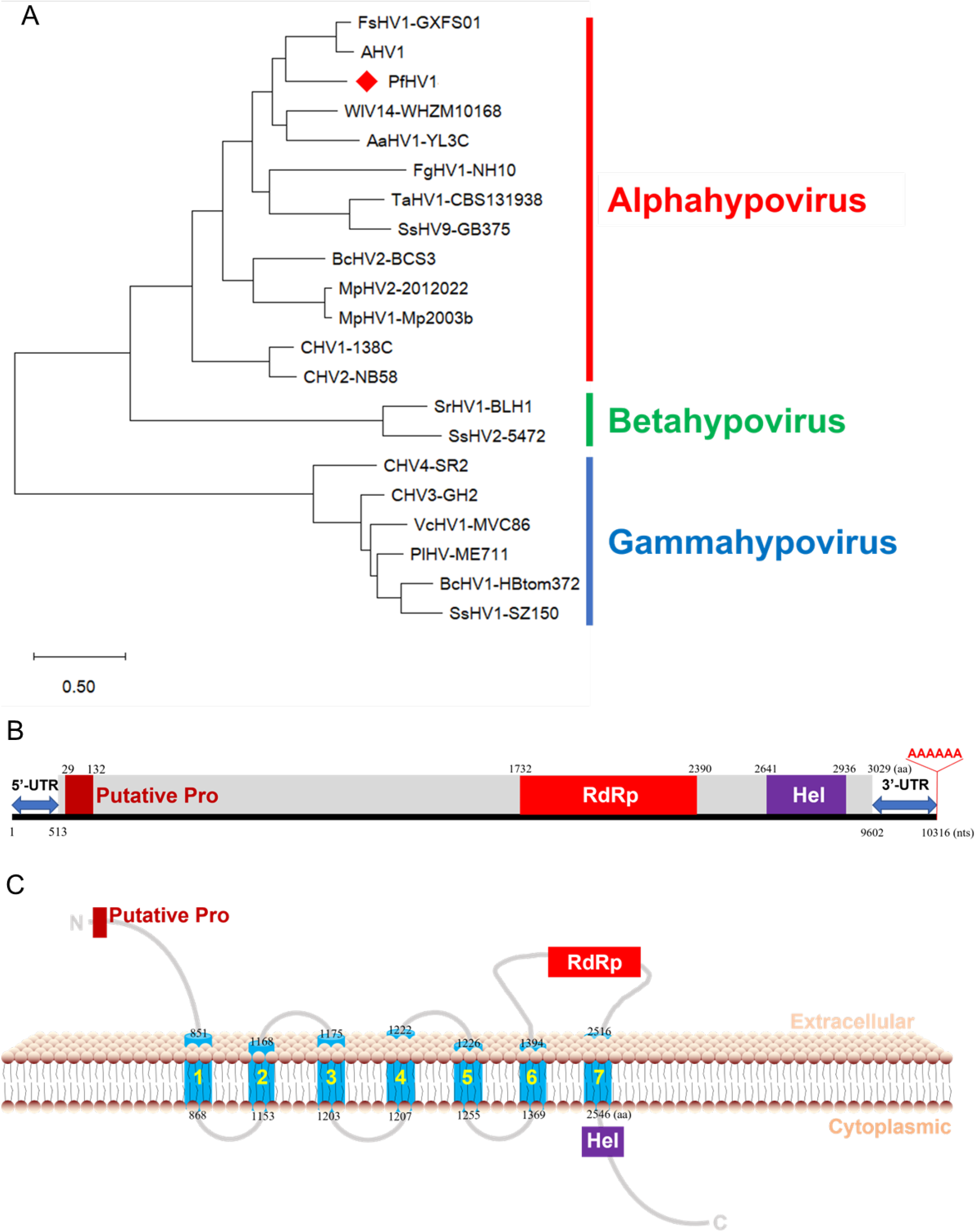
Phylogenetic analysis and genomic organization of Pestalotiopsis fici hypovirus1 (PfHV1). (A) A ML phylogenetic tree constructed based on the polyprotein sequences encoded by PfHV1 dsRNA1 open reading frame (ORF1) and those of known hypoviruses. (B) Genomic organization of the dsRNA1-encoding polyprotein (grey color). The conserved motifs for protease (Pro), RNA dependent RNA polyprotein (RdRp) and helicase (Hel) domain blocks on the polyprotein are marked in dark red, red and purple respectively, with the lengths corresponding to their aa sizes. The numbers under the line indicate the start and end positions of genome, 5′- and 3′-untranslated region (UTR), and the conserved domains. (C) Transmembrane domains (TMDs) predicted on the PfHV1 polyprotein. Blue cylinders indicate the TMDs fixed in the plasmalemma.

**Figure 3.**
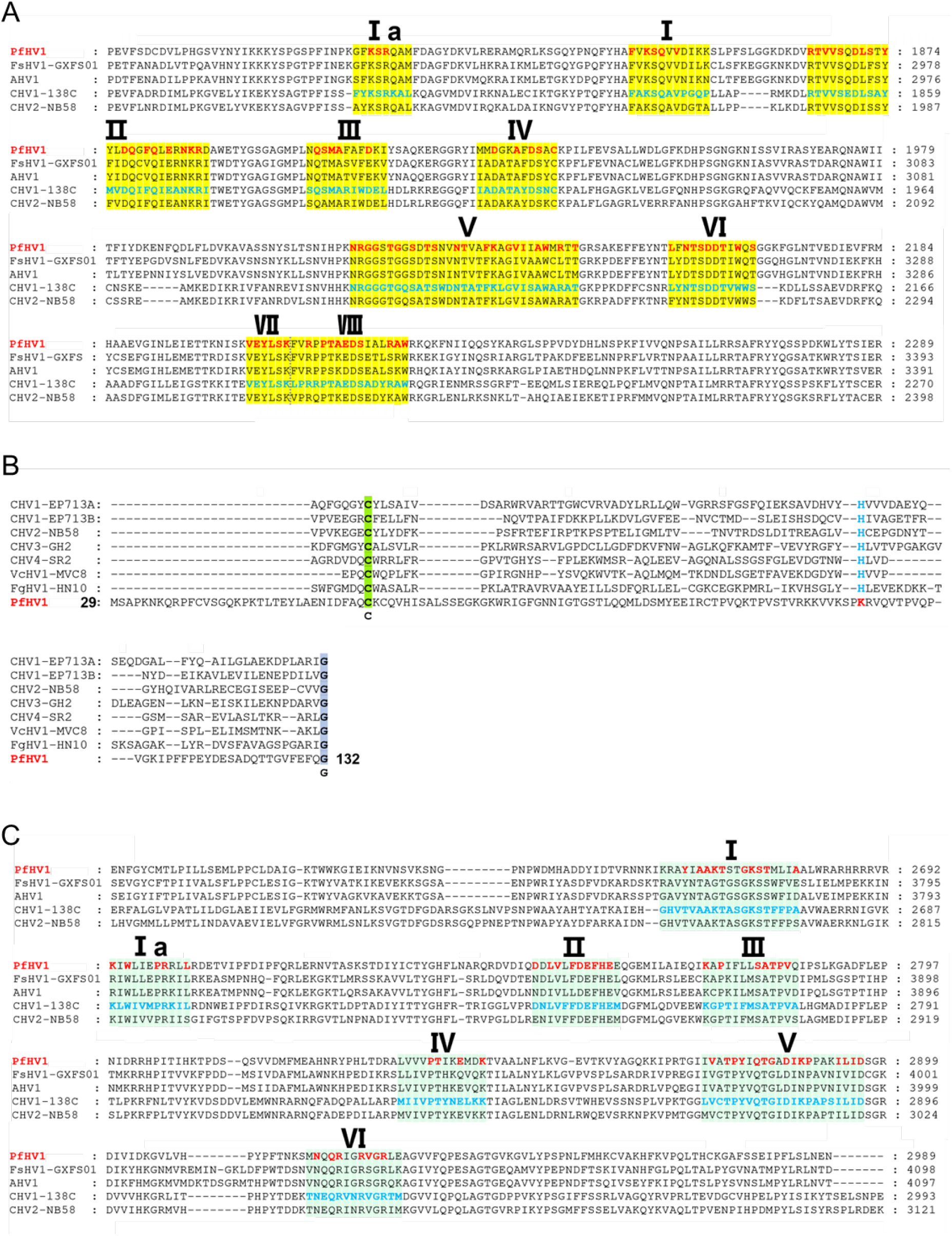
Alignment of RdRp, Pro and Hel with those of representative hypoviruses. (A to C) The conserved domains for RdRp, Pro, Hel. The conserved motifs or residues are highlighted.

BLASTp searches revealed that the polypeptide has the highest identity (55.13%) with the polyprotein (accession no. YP_009342443.1, coverage 96%, *E* value = 0) encoded by Wuhan insect virus 14 (WIV14), as well as high identities (50.25-55.10%, coverage 55–92%, *E* value = 0) to those of Fusarium sacchari hypovirus 1 (FsHV1; no. QIQ28422.1; Yao et al., 2020), Apis hypovirus 1-3 (no. UCR92523.1, UCR92524.1, UCR92525.1; direct submission), and Alternaria alternata hypovirus 1 (AaHV1; no. QFR36339.1; Li et al., 2019; Tab. S2). Phylogenetic analysis of the polyprotein sequence with those of all the known members of the family *Hypoviridae* illustrated that the sequence clustered with FsHV1 and AaHV1 in the newly proposed genus “Alphahypovirus” (3, 5). These results suggest that dsRNA1 is a genomic component of a novel hypovirus, and it is tentatively named Pestalotiopsis fici hypovirus 1 (PfHV1).

### dsRNA2 is a D-RNA of PfHV1

The complete nucleotide sequence of dsRNA2 is 5511 nts in length excluding the poly (A) tail, consisting of a single putative ORF (nt positions 510-4043), coding for a polyprotein of 1177 aa with an approximate molecular mass of 133.5 kDa. Alignment with dsRNA1 indicated that the total sequence of dsRNA2 is separately identical with dsRNA1 in five regions, i.e., nt position 4 to 3939, 5988 to 6225, 6512 to 6770, 6220 to 6508, and 6774 to 7562. Namely, dsRNA2 is a D-RNA generated from dsRNA1 by deletion three regions from nts 1 to 3, 3940 to 5987 and 7563 to 10316 (Figure 4A). Moreover, an inversion was observed in dsRNA2 sequence between nt position 4175 to 4433 and 4434 to 4722 as compared with dsRNA1 (Figure 4A).

**Figure 4.**
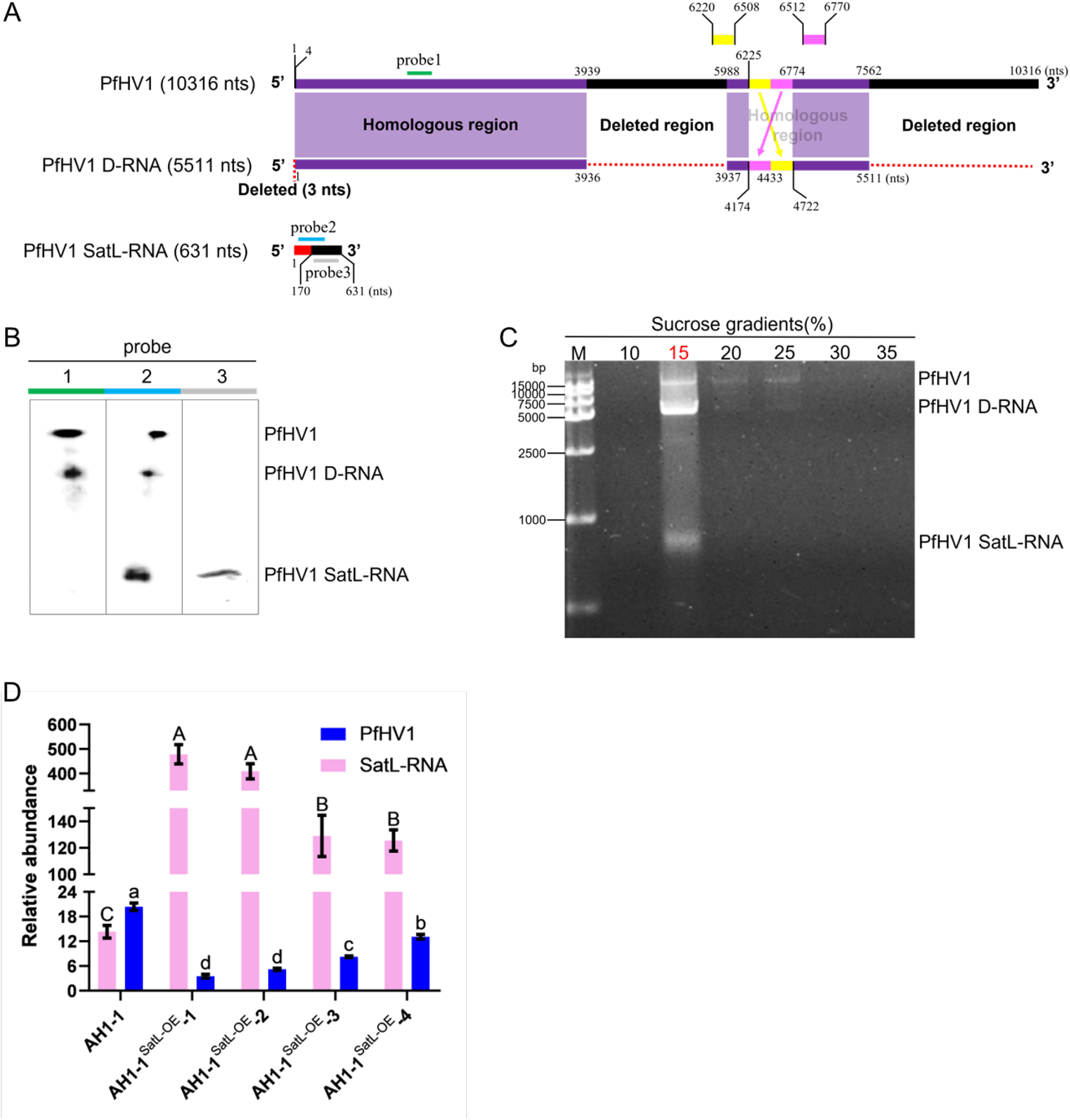
Identification of the defective RNA (D-RNA) and satellite-like dsRNA (SatL-dsRNA) of PfHV1. (A) Schematic diagram of the corresponding positions of dsRNA1, D-RNA, and SatL-dsRNA sequences. The purple and black blocks indicate the identical and deleted regions in dsRNA1 and D-RNA, respectively, and yellow and pink lines indicate the inversed regions in them. The red and black blocks indicate identical and exogenous regions in SatL-dsRNA and dsRNA1 or and D-RNA, respectively. (B) Northern blotting analysis of dsRNA1, D-RNA and SatL-dsRNA using riboprobe 1 to 3, whose corresponding positions in these dsRNAs are marked in green, blue, and grey, respectively. (C) Co-precipitation analysis of dsRNA1, D-RNA and Sat-RNA by ultra centrifugation in stepwise sucrose gradients (from 10% to 35% sucrose with 5% increments). (D) Relative expression analysis of dsRNA1 and SatL-RNA using RT-qPCR. Data are means±SEM (n = 3). Different letter indicates a significant difference at p < 0.05 (one-way ANOVA).

To further confirm that dsRNA2 as a D-RNA generated from dsRNA1, a digoxigenin (DIG)-labeled riboprobe (termed riboprobe 1) binding to the nt 1588 to 1934 of dsRNA1 was synthesized *in vitro* transcription and hybridization with the nucleic acid preparations of *P. fici* strain AH1-1. As expected, two hybridization signals were observed corresponding to the electrophoresis position of dsRNAs 1 and 2, further supporting the D-RNA nature of dsRNA2.

### dsRNA3 is a satellite-like dsRNA (SatL-dsRNA) of PfHV1

The complete nucleotide sequence of dsRNA3 is 631 bp in length without a poly (A) tail, and it is a non-coding RNA although it has a small putative ORF (nt positions 248-398) on its positive strand coding for a protein of 49 aa with an approximate molecular mass of 5.6 kDa, most likely it is not a *bona fide* protein due to so small size. Alignment of dsRNA3 with dsRNAs 1 and 2 indicated that their first 170 bp were 100% identical, excluding 3 nts deletion at 5′-termini of dsRNA 2, while the remaining sequence had no detectable similarity with both dsRNAs. To further confirm that dsRNA3 is identical with dsRNAs 1 and 2 at the 5′-termini while heterogenetic in the remaining, two DIG-labeled riboprobes (termed riboprobe 2 and 3) binding to the identical and heterogenetic regions of dsRNA1, respectively, were synthesized *in vitro* transcription and hybridization with the nucleic acid preparations of *P. fici* strain AH1-1. As riboprobe 2 was utilized, three hybridization signals were observed corresponding to the electrophoresis position of dsRNAs 1 to 3; while as riboprobe 3 was utilized, only one hybridization signal was observed corresponding to the electrophoresis position of dsRNA3 (Fig. 4B). Thus, the hybridization analysis supports the conclusion from the sequence alignment.

To further confirm that dsNRA3 is a SatL-dsRNA hosted by PfHV1, the presumed viral vesicles were purified by centrifugation in stepwise sucrose gradients (10–35% with 5% sucrose increments), and subjected to nucleic acids extraction. Agarose gel electrophoresis of the resulted nucleic acids showed that dsRNA3 together with the typical pattern of PfHV1 dsRNAs 1 and 2 was most recovered from the 15% fractions (Fig. 4C). These data suggest that dsRNAs 1 to 3 are encapsulated together, and it supports that dsNRA3 is a SatL-dsRNA of PfHV1.

### dsRNA3 negatively regulates the replication of PfHV1

To investigate whether the dsRNA3 affects the replication of PfHV1, the full-length of dsRNA3 cDNAs was engineered into an expression vector containing a hygromycin resistance marker, and transfected into the protoplasts prepared from the PfHV1-free strain AH1-1V^-^ of *P. fici*. With the resistance selection and fluorescence selection, three derivative subisolates AH1-1^SatL-OE^ transfected by dsRNA3-expression vector were obtained after being confirmed by RT-PCR detection of the RNA fragment (Fig.S2A), which were further dual-cultured with AH1-1 mycelia to generate the subisolates infected by PfHV1 in combination with dsRNA3-expression. Four hygromycin-resistant AH1-1 subisolates (termed AH1-1^SatL-OE^-1 to -4) derived from the dual-culture were randomly selected for nucleic extraction and analyzed on agarose gel, and revealing that they were all infected by dsRNAs 1 and 2 (Fig. 5C). Expression analysis of dsRNA1 together with dsRNA3 using RT-qPCR in the four resulting sub-isolates revealed that the expression levels of RNA3 were significantly up-regulated up to 8.77 to 33.39 folds in these transfected subisolates (Fig. 4D), whereas dsRNA1 were significantly down-regulated, up to 35.67 to 82.63 percent as compared with the parental strain AH1-1, in which, both dsRNAs had a similar expression level (Fig. 4D). These results suggest that the overexpression of the satellite-like RNA inhibits the replication of PfHV1.

**Figure 5.**
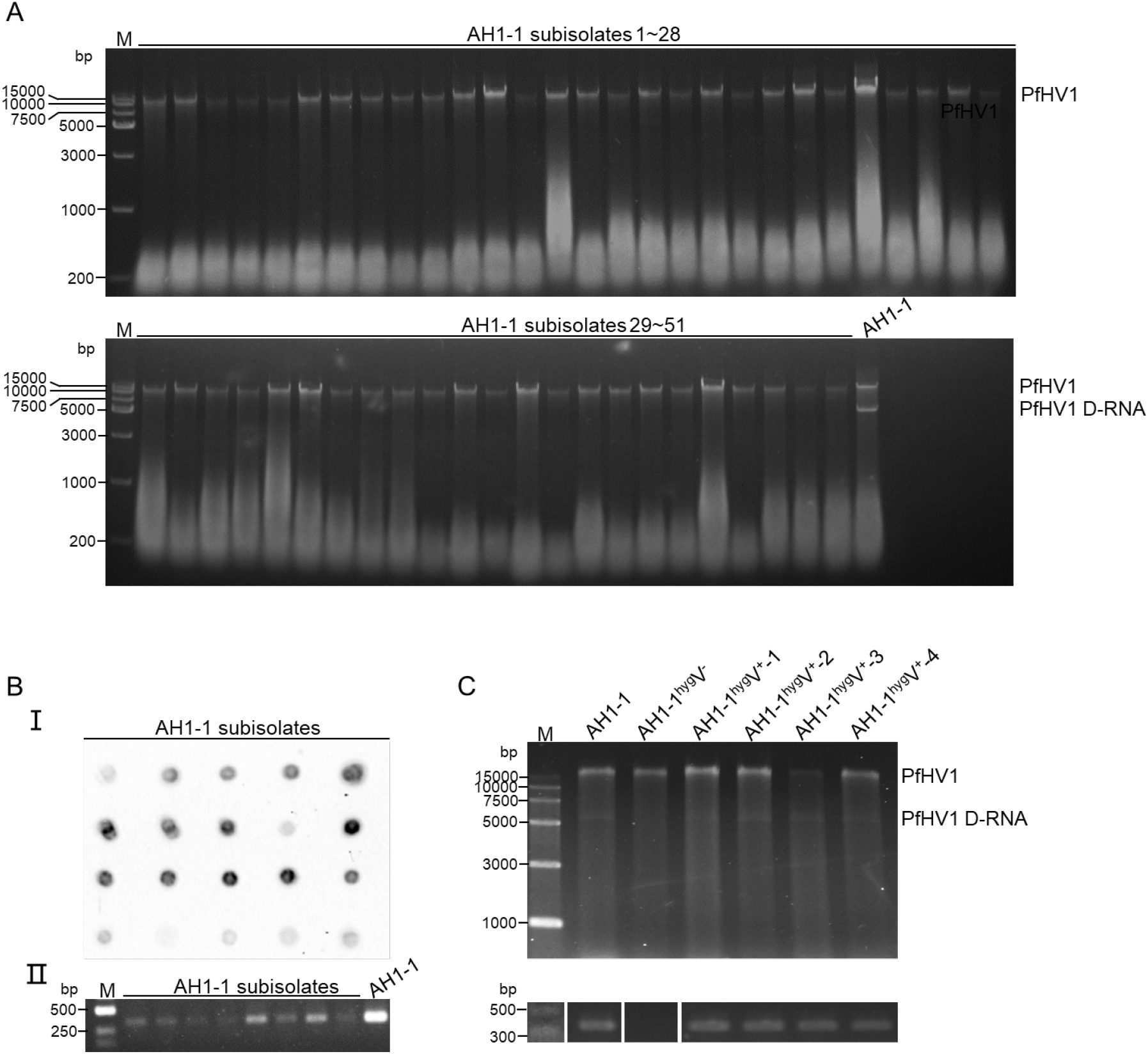
Electrophoresis and dot blotting analysis of horizontal and vertical transmission of PfHV1. Electrophoresis analysis of dsRNAs extracted from the conidium-generated subisolates of strain AH1-1 on 1% agarose gel after being treated by DNase I. (B) Dot blotting analysis with riboprobe 1 (I) and RT-PCR detection with primer pair PfHV1-1F/-1R (II, Table S1) of PfHV1 in some randomly selected conidium-generated subisolates. (C) Electrophoresis analysis on 1% agarose gel of dsRNA extraction and RT-PCR analysis with primer pair PfHV1-1F/-1R of PfHV1 in the subisolates (AH1-1^hyg^V^+^-1 to -4) of AH1-1^hyg^V^-^ (as a recipient strain) contact culture with AH1-1 (donor).

### PfHV1 dsRNAs are transmitted vertically and horizontally in different efficiency

To investigate potential vertical transmission of PfHV1 in *P. fici* strain AH1-1, individual conidia were isolated from mycelia and cultured on PDA. A total of 51 single subisolates were randomly selected and analyzed for the presence of PfHV1 dsRNAs. The experiments showed all (100%) of the subisolates were infected with PfHV1 dsRNA1 as detected by agarose gel analysis of their nucleic acids after treated by DNase I (Fig. 5A). This was further confirmed by dot blotting analysis of 25 randomly selected subisolates using riboprobe 1 and RT-PCR analysis of 8 subisolates using primers PfHV1-1F/1R for dsRNA 1, respectively (Fig.5B). It is worth noting that the RT-PCR products of PfHV1 dsRNA1 showed fewer contents for all the derived subisolates as compared with the parental strain AH1-1 (Fig. 5B) whereas dsRNAs 2 or 3 were not detected in all these conidium-generated subisolates, suggesting that they are not vertically transmitted via conidia.

To investigate horizontal transmission of PfHV1, PfHV1-infected AH1-1 (donor) and AH1-1^hyg^V^-^ (receptor) were dual-cultured, and four AH1-1V^-^ mycelium discs next to the contact area were randomly selected for RT-PCR analysis of the presence of PfHV1 dsRNAs. This experiment revealed that all the selected subisolates (nominated AH1-1^hyg^V^+^-1 to -4) were infected by PfHV1 dsRNAs 1 and 2, while not by dsRNA3 (Fig. 5C). Whereas as dual-culture of AH1-1 and PfHV1-free *Pestalotiopsis* sp. CJB-1, none of 12 isolates from the later were infected by PfHV1 dsRNAs 1 to 3.

These results suggest that PfHV1 dsRNA1 is efficiently transmitted vertically and horizontally in the host strain, while not easily to other *Pestalotiopsis* strains. dsRNA2 is efficiently transmitted horizontally but not vertically in the same host strain, whereas dsRNA3 could not be transmitted horizontally and vertically.

### PfHV1 confers no obvious effects on the biological traits of the host fungus

To check whether PfHV1 confers some biological effects on the host fungus, the morphologies and growth rates were accessed for strain AH1-1 (PfHV1^+^) and its cured subisolate AH1-1V^-^ (PfHV1^-^) after culture on PDA in parallel at 4 dpi, no obvious difference was observed in these aspects (Fig. 6A, B). Moreover, they induced similar sized lesions as being inoculated on wounded tea leaves (*C. sinensis* var. Fuyun no.6) (Fig. 6C). As compared with other strains free of PfHV1 including *Pestalotiopsis* sp. strains (AH1-1-14, TP-2-2W, JWX-3-1, and CJB-4-1), they also showed similar morphologies, growth rates, and virulence (Fig. 6D, E). These results suggest that PfHV1 confers no obvious effects on the morphologies and virulence of *Pestalotiopsis* fungi.

**Figure 6.**
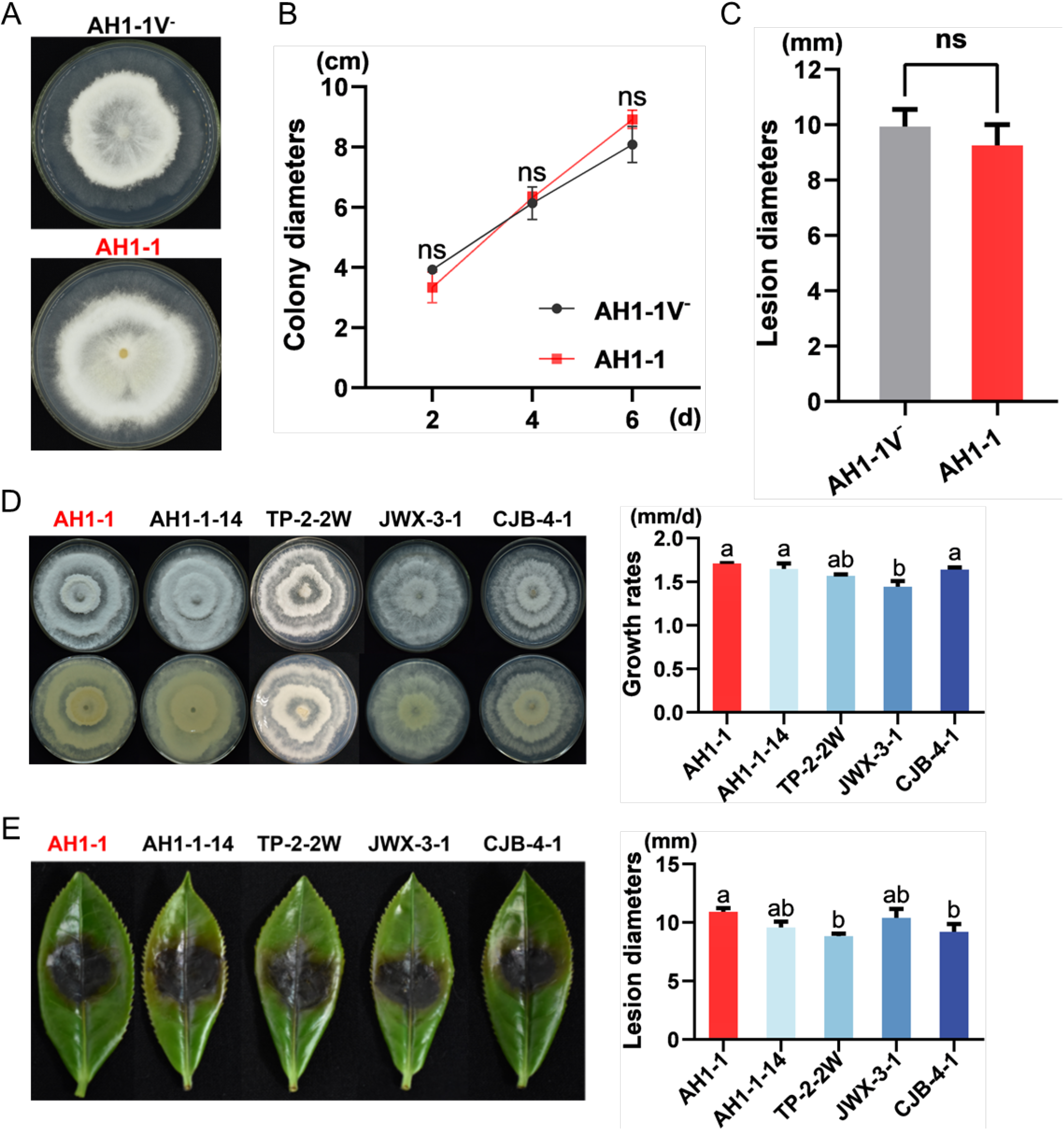
Assessment of the effects of PfHV1 infection on fungal growth and virulence. (A and B) Colonies and growth rates of AH1-1V^-^ (PfHV1^-^) and AH1-1 (PfHV1^+^) strains cultured on PDA medium for 6 days, respectively. The growth rates were measured at 2, 4, and 6 day, respectively. (C) The lesion lengths on tea leaves (*C. sinensis* var. Fuyun no.6) induced by inoculation with the mycelium discs of AH1-1V^-^ and AH1-1 strains at 4 day post inoculation (dpi). (D) Colonies and growth rates of AH1-1 and other *Pestalotiopsis* spp. strains on PDA medium cultured for 5 days. (E) The symptoms and lesion lengths caused by AH1-1 and other *Pestalotiopsis* spp. strains on tea leaves (*C. sinensis* var. Fuyun no.6), respectively. Data are means ± SEM (n = 6 or 3 for virulence and growth rate assessment, respectively). Different letter indicates a significant difference at p < 0.05 (one-way ANOVA). *** indicates a significant difference at p <0.01 (independent-samples t test), ns indicates no significant difference at p <0.05 (independent-samples t test).

## DISCUSSION

In this study, three dsRNA components were detected in *P. fici*, and their full-length sequences were determined and characterized, suggesting that they belong to a novel hypovirus, tentatively named PfHV1, under the newly proposed genus “Alphahypovirus” (3, 4, 25). Here, the dsRNA nature of PfHV1 genome was confirmed with enzymatic treatments, suggesting that PfHV1 genomic components survive as a dsRNA nature *in vivo*. As sucrose fraction ultra-centrifuge, the dsRNA components were co-precipitated in 15% sucrose fraction, and electrophoresed on agarose gel in the same position as compared with the ones extracted from the fungal mycelia, suggesting that its mature genome encapsulated in vesicles are still living with dsRNA nature. As hypoviruses are most closely related to potyviruses, a kind of positive single-stranded (+) RNA viruses infecting plants, while they have properties distinct from those of the potyviruses (23), we propose that PfHV1 together other hypoviruses may represent an intermediate stage in the evolution of a (+) ssRNA virus into a dsRNA virus. Other than CHV 1 to CHV4 which have been well studied in *Cryphonectria parasitica* (1), hypoviruses or hypo-like viruses have been largely discovered from other phytopathogenic fungi, including *Sclerotinia sclerotiorum* (10, 25–27) *Valsa ceratosperma* (3), *Fusarium graminearum* (7, 11), *Phomopsis longicolla* (28), *Macrophomina phaseolina* (29), *Botrytis cinerea* (6), and *Rosellinia necatrix* (30), and even in non-fungal eukaryotes (invertebrates) (31) revealed by large-scale meta-transcriptomic analysis. To our knowledge, this is the first report of hypovirus isolated from *Pestalotiopsis* fungus, and the second mycovirus besides of a chrysovirus infecting this fungal genus.

Genomic analysis of PfHV1 dsRNA1 suggests that it contains one ORF coding a polyprotein, which is somewhat different from other members in “Alphahypovirus”, since the group genomes typically harbor two ORFs (A and B), with two papain-like cysteine Pro encoded by ORF A and the leading regions of ORF B, respectively, as exemplified by CHV1 (23, 32, 33), while it is similar to the genomic organization of AaHV1 and those of genus “Betahypovirus” and “Gammahypovirus”. Moreover, a complete papain-like cysteine Pro motif was not detected in the polyprotein of PfHV1, whereas alignment with the polyproteins of other hypoviruses uncovered the presence of two conserved cysteine Pro core residues (cysteine, and glycine in Fig. 3B; 22) but lacked the histidine at its N-terminal region (29–132 aa region), conserved among hypo- and hypo-like viruses (5), suggesting that the cysteine Pro might be encoded by PfHV1 dsRNA1 to process the polyprotein, but it possess an obvious diversity and evolution at this region of PfHV1 genome. Moreover, the conserved domains of ORFs coding for the functional proteins including RdRp and Hel were detected in the polyprotein of PfHV1. Of these, RdRp is important for RNA viral replication and always conserved in polyproteins encoded by hypo- and hypo-like viruses (5, 24). While several aa were divergent in the conserved motifs as compared with the known members, e.g., in RdRp motif IV and VI. More importantly, a conserved SDD tripeptide was observed in RdRp motif IV, and it is different from most known (+) ssRNA viruses, in which the consensus tripeptide is GDD (34, 35), supporting our previous conclusion that PfHV1 may represent an intermediate stage in the evolution of a (+) ssRNA virus into a dsRNA virus. Several TMDs were predicted for PfHV1 polyprotein (Fig. 2C), and it is most likely that the cleaved proteins including Pro, RdRp and Hel associated with the vesicle membranes, and utilize the fungal *trans*-Golgi network for replication as predicted for CHV1 (36).

dsRNA 2 is determined as a defective component of dsRNA 1, which is very stable since it could be horizontally transmitted, as well as easily observed in other natural *Pestalotiopsis* spp. strains (data not shown), like the defective RNA 2 associated with CHV3 (37). However, dsRNA 2 could not be vertically transmitted together with dsRNA1, suggesting that is not a normal and necessary component of PfHV1 but rather a by-product of virus replication. It is worth noting that it is composed of two inversed sequences of considerately large size, suggesting that a deletion-junction process had happened after the transcription of dsRNA1, which is like a (+) ssRNA virus, in which the defective RNAs is maintenance of an ORF through a deletion-junction to maintain the translation of two different genes (38, 39). The coding capacity of the defective RNA and ability to be translated may affect accumulation (40, 41), since we have observed the PfHV1 dsRNA1 turned into an obviously lower titer after vertical transmission in the fungal progeny subisolates without the presence of the dsRNA2 components. It is unknown what viral factors are involved in generation and maintenance of the defective dsRNA, which is likely cleaved by dicer-like 2 (DCL2) and argonaute-like 2 (AGL2) proteins responsible for the antiviral RNA silencing pathways in a filamentous fungus (9, 42).

dsRNA3 segment is determined to be a SatL-dsRNA of PfHV1 due to the following reasons: 1) it was co-precipitated with dsRNAs 1 and 2 in the same sucrose fraction of 15% by ultra-centrifuge, suggesting that it is encapsulated together with PfHV1 genomic dsRNAs; 2) it codes no structural proteins or RdRp proteins, suggesting its replication depends on the helper virus; 3) no dsRNA3 component was solely observed in any subisolates absent of dsRNA1 after vertical or horizontal transmission. Until now, three satellite or SatL-dsRNAs have been observed in association with hypoviruses, i.e., dsRNA4 (937 bp in size) and dsRNA3 (dimeric dsRNA4 linking by a poly (A)) of CHV3 in *C. parasitica* and S-dsRNA (3.6 kb) of SsHV1 in *S. sclerotiorum*, both of which encode authentic proteins and contain a poly (A) tail (10, 37). Of these, CHV3 dsRNA-4 was confirmed to encode a small polypeptide of 9.4 kDa with an *in vitro* translation experiment (37, 43); S-dsRNA (3.6 kb) of SsHV1 was proposed to encode a protein of 639 aa with an approximate molecular mass of 71 kDa (10). PfHV1 dsRNA3 shares no detectable identities with the both satellite (or satellite-like) dsRNAs, contains no a poly (A) tail, and most likely encodes no proteins, suggesting that dsRNA3 is a completely different SatL-dsRNA as compared the ones of CHV3 and SsHV1, and belongs to a new class of satellite nucleic acids. Generally, satellites are distinct from their helper virus with a nucleotide sequence that is substantially different from that of their helper virus, generally having very little or no sequence similarity with helper viruses, encapsidated by the CP of helper viruses (44). Here, dsRNA3 should be encapsulated in the replication vesicles, composed of host-derived lipids (45), instead of CP by the helper virus PfHV1 since the latter has no structural proteins; moreover, dsRNA3 shares a substantially identical sequence (170 bp in size) with the helper virus genome, suggest that it partially originated from its helper virus and is distinct from a typical satellite nucleic acid. Therefore, PfHV1 dsRNA3 represents a novel class of satellite nucleic acids unreported before, unlike the dsRNA satellites in association with *Totiviridae* and *Partitiviridae*. We propose these satellite nucleic acids are classified into two classes including those coding dsRNAs with a poly (A) tail (like CHV3 dsRNAs 3 and 4) and those no-coding dsRNAs without a poly (A) tail (like PfHV1 dsRNA3). Additionally, we suggest to broaden the definition of satellites to accommodate these satellite dsRNAs by including a special type of satellite nucleic acid that has substantial sequence homology with the host viral genome without encapsidation in a coat protein.

Satellite dsRNAs have been described in association with several mycoviruses in filamentous fungi and yeast (46–48), and it has been proved that the amplification used viral RdRp (12, 49). Of them, the satellite dsRNAs associated with *Totiviridae* encode a secreted preprotoxin that is lethal to sensitive cells (virus-free or containing helper virus only), and imparts self-protection against the secreted toxin and confers ecological advantage by killing competing virus- or satellite-free fungi; the satellite dsRNA associated with SsHV1 most likely enhances the hypovirulence traits of *S. sclerotiorum*. However, the biological functions of other satellite dsRNAs remain largely unknown. Here, as compared with the dsRNA3-infected and -free strains of *P. fici* isolates, i.e., strain AH1-1 subisolates derived from horizontal transmission and those from vertical transmission, no obvious morphologies, growth rates or virulence were observed, suggesting that the SatL-dsRNA has no impact on these aspects. However, as the satellite-like RNA was overexpressed, it inhibits the replication of PfHV1, suggesting that dsRNA3 negatively regulates the replication of PfHV1. These results suggest that the SatL-dsRNAs harbored by hypoviruses have a similar biological trait to the satellite RNAs associated with animal and plant viruses. E.g., amplification of satellites by viral RdRp may down-regulate synthesis of viral RNAs and expression of their products in the research of Trichomonas vaginalis virus (49); cucumber mosaic virus (CMV) satellite RNA reduced the yield of accumulated helper virus (50, 51). Based on the *in vitro* competition assay, the reduction was proposed to be due to the competition of satellite RNA for limited amount of viral replicase with CMV gRNAs (52). Furthermore, a three-way branched secondary structure was identified in the satellite RNA, and was indispensable for the helper virus inhibition (53). Similarly, a conserved apical hairpin stem-loop (AHSL) structure was identified in 5′-UTR satellite RNA of bamboo mosaic virus (BaMV) as a potent determinant of the down-regulation of helper virus replication (54). Since the SatL-dsRNA encodes no functional proteins, we conclude that it inhibits the replication of helper virus most likely with similar mechanism to the CMV and BaMV satellite RNAs.

In summary, to gain insight into mycoviruses in *P. fici*, belonging to an important fungus genus related to agriculture, industry and medicine, a novel hypovirus from *P. fici* together with its D-RNA and a SatL-dsRNA, were identified and characterized. To our knowledge, PfHV1 represents the first hypovirus and the second mycovirus isolated from *Pestalotiopsis* spp., and its SatL-dsRNA represents a novel class of satellite nucleic acids, which should contribute to our better understanding of the taxonomy, evolution, molecular and biological traits of mycoviruses, as well as the related satellites.

## Data availability statement

The complete nucleotide sequence of the PfHV1 genome was deposited in the GenBank database under accession numbers OP441373 to OP441375.

## Author contributions

W. X. designed the investigation, provided project supervision and funding support, and improved the manuscript. Z. H. wrote the initial manuscript, and conducted the horizontal transmission, transfection, expression analysis, partial hybridization, and partial pathogenicity assays, J. L. conducted genomic cloning, enzymatic treatment, vertical transmission, partial hybridization, and partial pathogenicity assays, D. N. and H. W. were involved in project supervision and funding acquisition, and improved the manuscript.

## ACKNOWLEDGEMENTS

This work was financially supported by the National Key Research and Development Program of China (No. 2021YFD1000400), National Natural Science Foundation of China (No. 31872014 and 32172475), Enshi Tujia and Miao Autonomous Prefecture Bureau of Science and Technology (No. XYJ2021000083), and Hubei Hongshan Laboratory to W.X.

## Figure legends

**Table S1.**
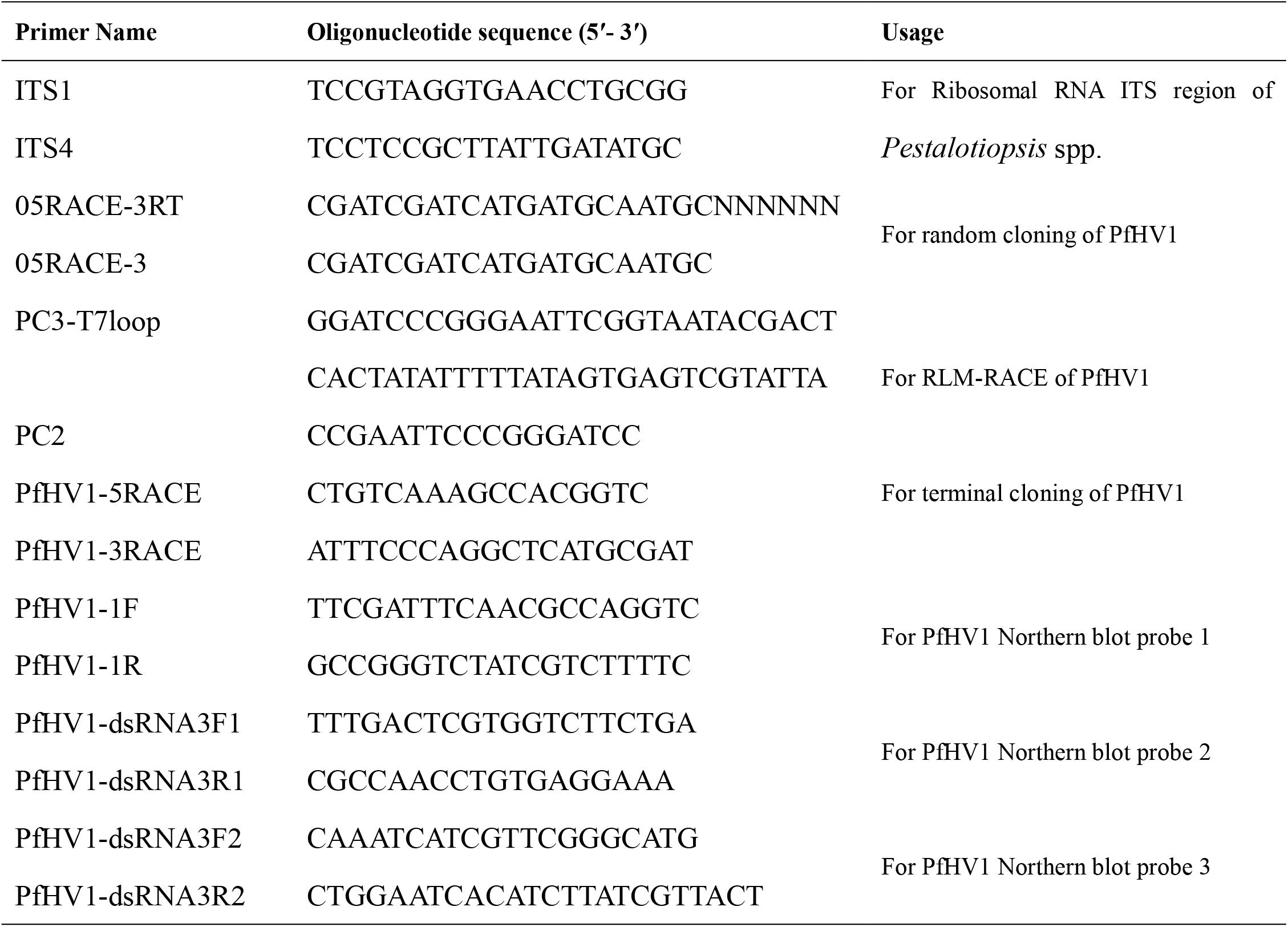
Primers used in this study.

**Table S2.** Blastp searches of dsRNA1-coding proteins with those deposited in NCBI.

| Scientific Name | Max<br>Score | Total<br>Score | Query<br>Cover | E value | Per. Ident | Acc.<br>Len | Accession |
| --- | --- | --- | --- | --- | --- | --- | --- |
| Fusarium sacchari hypovirus 1 | 2510 | 2703 | 87% | 0.0 | 55.10% | 4257 | QIQ28422.1 |
| Apis hypovirus 1 | 2483 | 2717 | 87% | 0.0 | 54.00% | 4256 | UCR92523.1 |
| Wuhan insect virus 14 | 2482 | 2804 | 96% | 0.0 | 55.13% | 3818 | YP_009342443.1 |
| Apis hypovirus 3 | 2416 | 2663 | 87% | 0.0 | 54.74% | 3383 | UCR92525.1 |
| Apis hypovirus 2 | 2409 | 2635 | 92% | 0.0 | 54.45% | 3525 | UCR92524.1 |
| Alternaria alternata<br>hypovirus 1 | 2241 | 2414 | 88% | 0.0 | 50.25% | 4228 | QFR36339.1 |
| Trichoderma asperellum<br>hypovirus 1 | 1814 | 1814 | 70% | 0.0 | 44.18% | 4176 | AZT88614.1 |
| Botrytis cinerea hypovirus 2 | 1773 | 1773 | 71% | 0.0 | 42.58% | 4199 | QJT73706.1 |
| Macrophomina phaseolina<br>hypovirus 2 | 1725 | 1931 | 90% | 0.0 | 42.67% | 4111 | QOE55583.1 |
| Sclerotinia sclerotiorum<br>hypovirus 9 | 1697 | 1697 | 70% | 0.0 | 41.43% | 4196 | UCR95333.1 |

**Figure S1.**
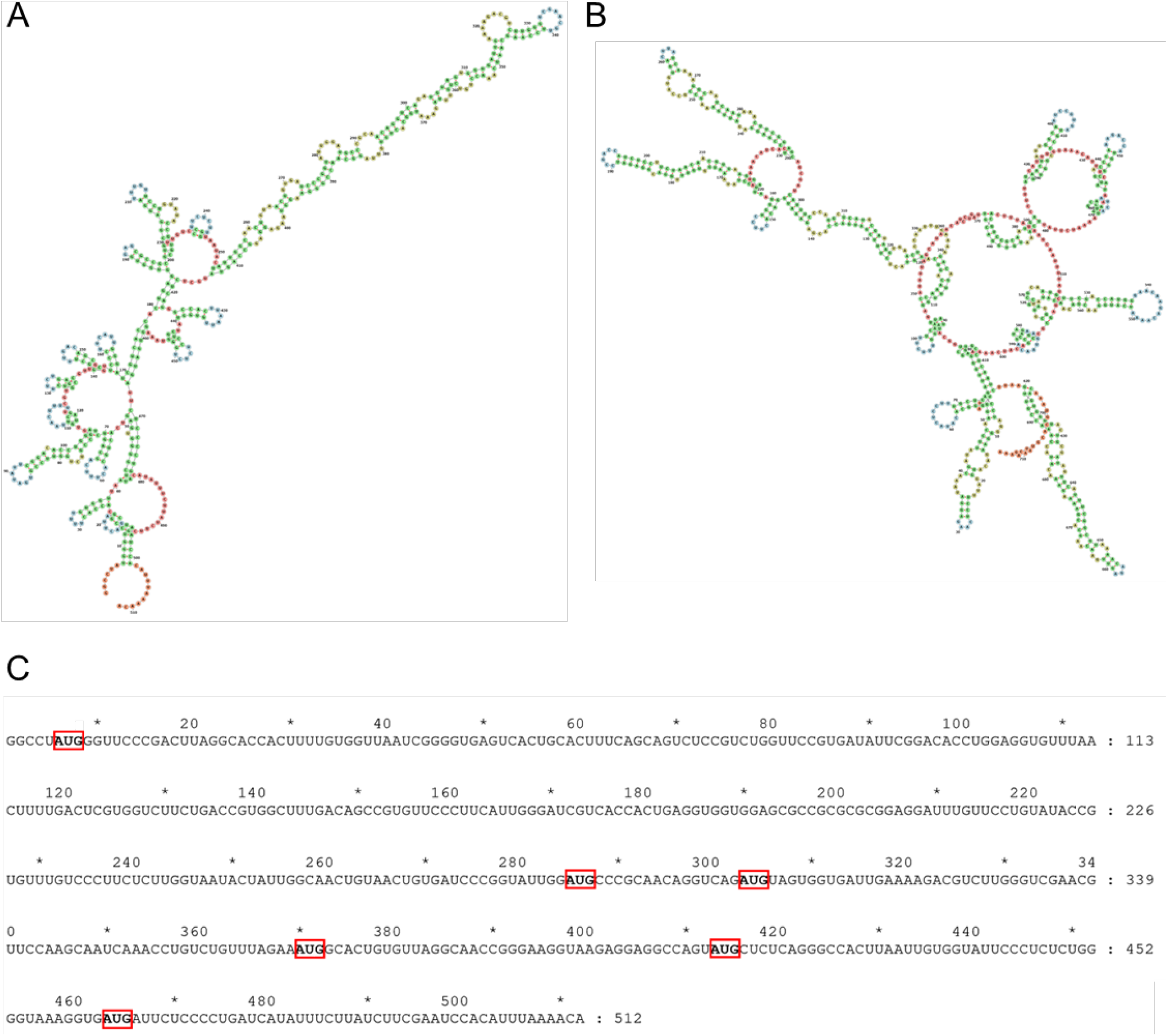
Predicted secondary structures of both UTR of PfHV1. (A and B) The secondary structures of 5′- and 3′-UTR of PfHV1 were predicted in RNAfold, respectively. (C) Six AUGs in the 5′-UTR of PfHV1 were found.

**Figure S2.**
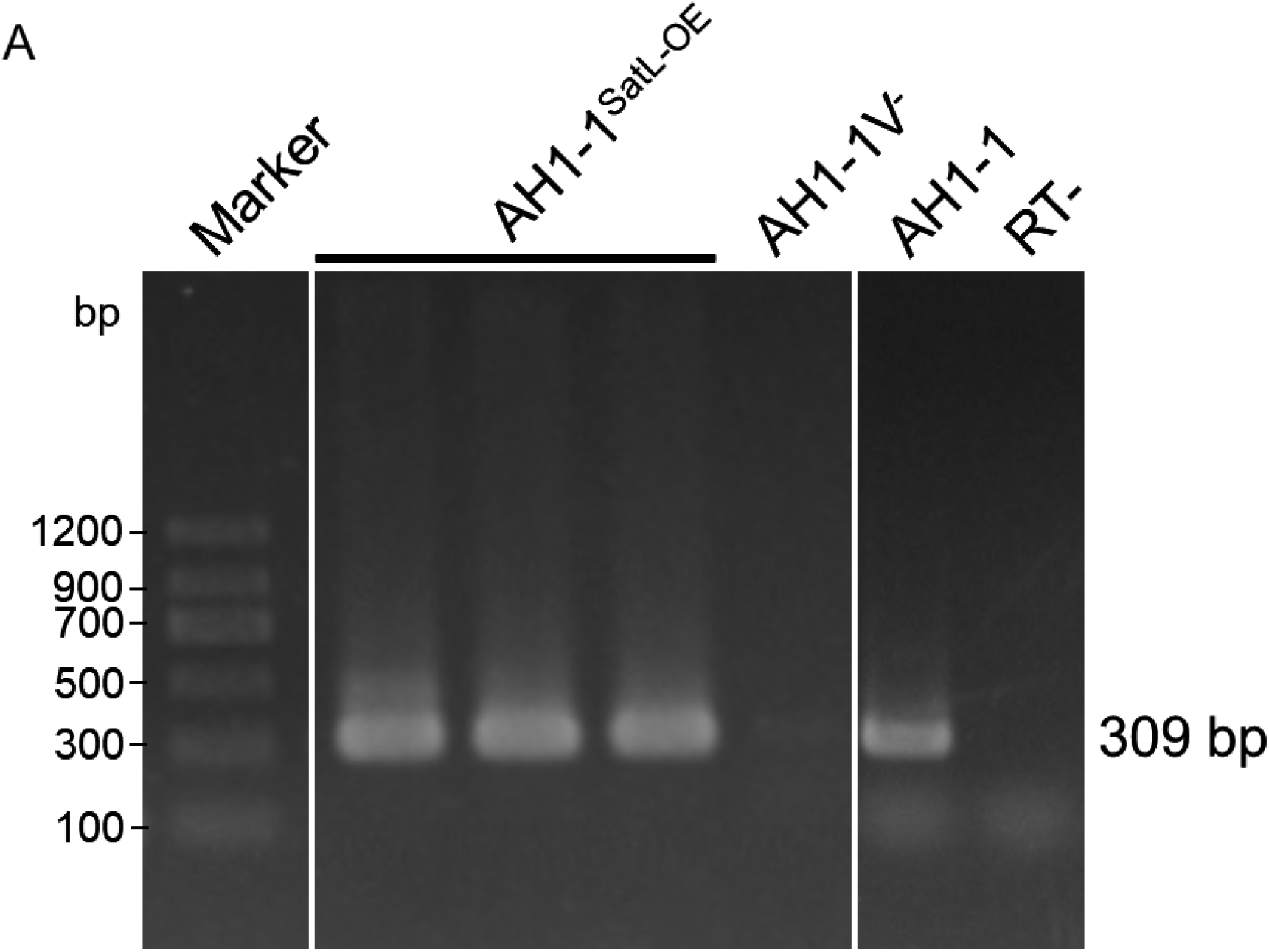
Identification of Sat-RNA in the protoplast-generated subisolates of AH1-1_SatL-OE_ after being transfected with an over-expressed vector. The over-expressed strain was generated by inserting the full-length of cDNAs of SatL-RNA strains into the *Pst* I cloning site of vector. The expression of Sat-RNA in the transfectants (AH1-1^SatL-OE^) was analyzed using RT-PCR with primers PfHV1-dsRNA3F2 and PfHV1-dsRNA3R2, with AH1-1 and AH1-1V^-^ involved as positive and negative controls, respectively. Lines2-4, strains derived from AH1-1 with SatL-RNA overexpression, AH1-1^SatL-OE^; Line5, PfHV1^-^ strain, AH1-1V^-^; Line6, PfHV1^+^ strain, AH1-1; Line7, blank control for reverse transcription.

## REFERENCES

1. Suzuki N, Ghabrial SA, Kim K-H, Pearson M, Marzano S-YL, Yaegashi H, Xie J, Guo L, Kondo H, Koloniuk I, Hillman BI, ICTV Report Consortium. 2018. ICTV virus taxonomy profile: Hypoviridae. J Gen Virol 99:615–616.

2. Hillman BI, Suzuki N. 2004. Viruses of the chestnut blight fungus, Cryphonectria parasitica. Adv Virus Res 63:423–472

3. Yaegashi H, Kanematsu S, Ito T. 2012. Molecular characterization of a new hypovirus infecting a phytopathogenic fungus, Valsa ceratosperma. Virus Res 165:143–150.

4. Khalifa ME, Pearson MN. 2014. Characterisation of a novel hypovirus from Sclerotinia sclerotiorum potentially representing a new genus within the Hypoviridae. Virology 464–465:441–449.

5. Li H, Bian R, Liu Q, Yang L, Pang T, Salaipeth L, Andika IB, Kondo H, Sun L. 2019. Identification of a novel hypovirulence-Inducing hypovirus from Alternaria alternata. Front Microbiol 10:1076.

6. Hao F, Ding T, Wu M, Zhang J, Yang L, Chen W, Li G. 2018. Two novel hypovirulence-associated mycoviruses in the phytopathogenic fungus Botrytis cinerea: molecular characterization and suppression of infection cushion formation. Viruses 10: 254.

7. Wang S, Kondo H, Liu L, Guo L, Qiu D. 2013. A novel virus in the family Hypoviridae from the plant pathogenic fungus Fusarium graminearum. Virus Res 174:69–77.

8. Shapira R, Choi GH, Hillman BI, Nuss DL. 1991. The contribution of defective RNAs to the complexity of viral-encoded double-stranded RNA populations present in hypovirulent strains of the chestnut blight fungus Cryphonectria parasitica. EMBO J 10:741–746.

9. Zhang X, Nuss DL. 2008. A host dicer is required for defective viral RNA production and recombinant virus vector RNA instability for a positive sense RNA virus. Proc Natl Acad Sci U S A 105:16749–16754.

10. Xie J, Xiao X, Fu Y, Liu H, Cheng J, Ghabrial SA, Li G, Jiang D. 2011. A novel mycovirus closely related to hypoviruses that infects the plant pathogenic fungus Sclerotinia sclerotiorum. Virology 418:49–56.

11. Li P, Zhang H, Chen X, Qiu D, Guo L. 2015. Molecular characterization of a novel hypovirus from the plant pathogenic fungus Fusarium graminearum. Virology 481:151–160.

12. Gnanasekaran P, Chakraborty S. 2018. Biology of viral satellites and their role in pathogenesis. Curr Opin Virol 33:96–105.

13. Zhou L, Li X, Kotta-Loizou I, Dong K, Li S, Ni D, Hong N, Wang G, Xu W. 2021. A mycovirus modulates the endophytic and pathogenic traits of a plant associated fungus. ISME J 15:1893–1906.

14. Wang L, He Y, Kang Y, Hong N, Farooq ABU, Wang G, Xu W. 2013. Virulence determination and molescular features of peach latent mosaic viroid isolates derived from phenotypically different peach leaves: a nucleotide polymorphism in L11 contributes to symptom alteration. Virus Res 177:171–178.

15. Maharachchikumbura SSN, Hyde KD, Groenewald JZ, Xu J, Crous PW. 2014. Pestalotiopsis revisited. Stud Mycol 79:121–186.

16. Nuss DL. 2005. Hypovirulence: mycoviruses at the fungal–plant interface. Nat Rev Microbiol 3:632–642.

17. Wang ZH, Zhao ZX, Hong N, Ni D, Cai L, Xu WX, Xiao YN. 2017. Characterization of causal agents of a novel disease inducing brown-black spots on tender tea leaves in China. Plant Dis 101:1802–1811.

18. Jia H, Dong K, Zhou L, Wang G, Hong N, Jiang D, Xu W. 2017. A dsRNA virus with filamentous viral particles. Nat Commun 8:168.

19. Zhai L, Xiang J, Zhang M, Fu M, Yang Z, Hong N, Wang G. 2016. Characterization of a novel double-stranded RNA mycovirus conferring hypovirulence from the phytopathogenic fungus Botryosphaeria dothidea. Virology 493:75–85.

20. Liu H, Fu Y, Jiang D, Li G, Xie J, Peng Y, Yi X, Ghabrial SA. 2009. A novel mycovirus that is related to the human pathogen Hepatitis E virus and rubi-like viruses. J Virol 83:1981–1991.

21. Tamura K, Stecher G, Kumar S. 2021. MEGA11 Molecular Evolutionary Genetics Analysis Version 11. Mol Biol Evol 38:3022–3027.

22. Pallás v, Sánchez-Navarro Ja, Kinard GR, Di Serio F. 2017. Chapter 35 - Molecular hybridization techniques for detecting and studying viroids, p. 369–379. In Hadidi, A, Flores, R, Randles, JW, Palukaitis, P (eds.), Viroids and satellites. Academic Press, Boston.

23. Koonin EV, Choi GH, Nuss DL, Shapira R, Carrington JC. 1991. Evidence for common ancestry of a chestnut blight hypovirulence-associated double-stranded RNA and a group of positive-strand RNA plant viruses. Proc Natl Acad Sci U S A 88:10647–10651.

24. Yao Z, Zou C, Peng N, Zhu Y, Bao Y, Zhou Q, Wu Q, Chen B, Zhang M. 2020. Virome identification and characterization of Fusarium sacchari and F. andiyazi: causative agents of pokkah boeng disease in sugarcane. Front Microbiol 11:240.

25. Hu Z, Wu S, Cheng J, Fu Y, Jiang D, Xie J. 2014. Molecular characterization of two positive-strand RNA viruses co-infecting a hypovirulent strain of Sclerotinia sclerotiorum. Virology 464–465:450–459.

26. Khalifa ME, Pearson MN. 2014. Characterisation of a novel hypovirus from Sclerotinia sclerotiorum potentially representing a new genus within the Hypoviridae. Virology 464–465:441–449.

27. Marzano S-YL, Hobbs HA, Nelson BD, Hartman GL, Eastburn DM, McCoppin NK, Domier LL. 2015. Transfection of Sclerotinia sclerotiorum with in vitro transcripts of a naturally occurring interspecific recombinant of Sclerotinia sclerotiorum hypovirus 2 significantly reduces virulence of the fungus. J Virol 89:5060–5071.

28. Koloniuk I, El-Habbak MH, Petrzik K, Ghabrial SA. 2014. Complete genome sequence of a novel hypovirus infecting Phomopsis longicolla. Arch Virol 159:1861–1863.

29. Marzano S-YL, Domier LL. 2016. Novel mycoviruses discovered from metatranscriptomics survey of soybean phyllosphere phytobiomes. Virus Res 213:332–342.

30. Arjona-Lopez JM, Telengech P, Jamal A, Hisano S, Kondo H, Yelin MD, Arjona-Girona I, Kanematsu S, Lopez-Herrera CJ, Suzuki N. 2018. Novel, diverse RNA viruses from Mediterranean isolates of the phytopathogenic fungus, Rosellinia necatrix: insights into evolutionary biology of fungal viruses. Environ Microbiol 20:1464–1483.

31. Shi M, Lin X-D, Tian J-H, Chen L-J, Chen X, Li C-X, Qin X-C, Li J, Cao J-P, Eden J-S, Buchmann J, Wang W, Xu J, Holmes EC, Zhang Y-Z. 2016. Redefining the invertebrate RNA virosphere. Nature 540:539–543.

32. Choi GH, Pawlyk DM, Nuss DL. 1991. The autocatalytic protease p29 encoded by a hypovirulence-associated virus of the chestnut blight fungus resembles the potyvirus-encoded protease HC-Pro. Virology 183:747–752.

33. Shapira R, Nuss DL. 1991. Gene expression by a hypovirulence-associated virus of the chestnut blight fungus involves two papain-like protease activities. Essential residues and cleavage site requirements for p48 autoproteolysis. J Biol Chem 266:19419–19425.

34. Poch O, Blumberg BM, Bougueleret L, Tordo N. 1990. Sequence comparison of five polymerases (L proteins) of unsegmented negative-strand RNA viruses: theoretical assignment of functional domains. J Gen Virol 71 (Pt 5):1153–1162.

35. Xu X, Liu Y, Weiss S, Arnold E, Sarafianos SG, Ding J. 2003. Molecular model of SARS coronavirus polymerase: implications for biochemical functions and drug design. Nucleic Acids Res 31:7117–7130.

36. Jacob-Wilk D, Turina M, Van Alfen NK. 2006. Mycovirus cryphonectria hypovirus 1 elements cofractionate with trans-Golgi network membranes of the fungal host Cryphonectria parasitica. J Virol 80:6588–6596.

37. Hillman BI, Foglia R, Yuan W. 2000. Satellite and defective RNAs of Cryphonectria hypovirus 3-Grand Haven 2, a virus species in the family Hypoviridae with a single open reading frame. Virology 276:181–189.

38. Romero J, Huang Q, Pogany J, Bujarski JJ. 1993. Characterization of defective interfering RNA components that increase symptom severity of broad bean mottle virus infections. Virology 194:576–584.

39. White KA, Bancroft JB, Mackie GA. 1991. Defective RNAs of clover yellow mosaic virus encode nonstructural/coat protein fusion products. Virology 183:479–486.

40. Pogany J, Romero J, Huang Q, Sgro JY, Shang H, Bujarski JJ. 1995. De novo generation of defective interfering-like RNAs in broad bean mottle bromovirus. Virology 212:574–586.

41. White KA, Bancroft JB, Mackie GA. 1992. Coding capacity determines in vivo accumulation of a defective RNA of clover yellow mosaic virus. J Virol 66:3069–3076.

42. Chiba S, Suzuki N. 2015. Highly activated RNA silencing via strong induction of dicer by one virus can interfere with the replication of an unrelated virus. Proc Natl Acad Sci U S A 112:E4911–4918.

43. Yuan W, Hillman BI. 2001. In vitro translational analysis of genomic, defective, and satellite RNAs of Cryphonectria hypovirus 3-GH2. Virology 281:117–123.

44. Bruening, G. 2002. Virus-dependent RNA agents, p 1170–1177. In Maloy O, Murray T (ed), Encyclopedia of plant pathology, vol 2. Wiley, New York, NY.

45. Newhouse JR, MacDonald WL, Hoch HC. 1990. Virus-like particles in hyphae and conidia of European hypovirulent (dsRNA-containing) strains of Cryphonectria parasitica. Can J Bot 68:90–101.

46. Deng F, Boland GJ. 2004. A satellite RNA of Ophiostoma novo-ulmi mitovirus 3a in hypovirulent isolates of Sclerotinia homoeocarpa. Phytopathology 94:917–923.

47. Wickner RB. 1996. Double-stranded RNA viruses of Saccharomyces cerevisiae. Microbiol Rev 60:250–265.

48. Romanos MA, Buck KW, Rawlinson CJ. 1981. A satellite double-stranded RNA in a virus from Gaeumannomyces graminis. J GEN VIROL 57.375–385

49. Khoshnan A, Alderete JF. 1995. Characterization of double-stranded RNA satellites associated with the Trichomonas vaginalis virus. J Virol 69:6892–6897.

50. Gal-On A, Kaplan I, Palukaitis P. 1995. Differential effects of satellite RNA on the accumulation of cucumber mosaic virus RNAs and their encoded proteins in tobacco vs zucchini squash with two strains of CMV helper virus. Virology 208:58–66.

51. Liao Q, Zhu L, Du Z, Zeng R, Peng J, Chen J. 2007. Satellite RNA-mediated reduction of cucumber mosaic virus genomic RNAs accumulation in Nicotiana tabacum. Acta Biochim Biophys Sin (Shanghai) 39:217–223.

52. Wu G, Kaper JM. 1995. Competition of viral and satellite RNAs of cucumber mosaic virus for replication in vitro by viral RNA-dependent RNA polymerase. Res Virol 146:61–67.

53. He L, Wang Q, Gu Z, Liao Q, Palukaitis P, Du Z. 2019. A conserved RNA structure is essential for a satellite RNA-mediated inhibition of helper virus accumulation. Nucleic Acids Res 47:8255–8271.

54. Hsu Y-H, Chen H-C, Cheng J, Annamali P, Lin B-Y, Wu C-T, Yeh W-B, Lin N-S. 2006. Crucial role of the 5′ conserved structure of bamboo mosaic virus satellite RNA in downregulation of helper viral RNA replication. J Virol 80:2566–2574.

